# Transcription factor FfmA interacts both physically and genetically with AtrR to properly regulate gene expression in the fungus *Aspergillus fumigatus*

**DOI:** 10.1101/2023.06.06.543935

**Authors:** Sanjoy Paul, Mark A. Stamnes, W. Scott Moye-Rowley

**Affiliations:** From: Department of Molecular Physiology and Biophysics, Carver College of Medicine, University of Iowa, Iowa City, IA. 52242 USA

## Abstract

Transcriptional regulation of azole resistance in the filamentous fungus *Aspergillus fumigatus* is a key step in development of this problematic clinical phenotype. We and others have previously described a C2H2-containing transcription factor called FfmA that is required for normal levels of voriconazole susceptibility and expression of an ATP-binding cassette transporter gene called *abcG1*. Null alleles of *ffmA* exhibit a strongly compromised growth rate even in the absence of any external stress. Here we employ an acutely repressible doxycycline-off form of *ffmA* to rapidly deplete FfmA protein from the cell. Using this approach, we carried out RNA-seq analyses to probe the transcriptome of *A. fumigatus* cells that have been deprived of normal FfmA levels. We found that 2000 genes were differentially expressed upon depletion of FfmA, consistent with the wide-ranging effect of this factor on gene regulation. Chromatin immunoprecipitation coupled with high throughput DNA sequencing analysis (ChIP-seq) identified 530 genes that were bound by FfmA using two different antibodies for immunoprecipitation. More than 300 of these genes were also bound by AtrR demonstrating the striking regulatory overlap with FfmA. However, while AtrR is clearly an upstream activation protein with clear sequence specificity, our data suggest that FfmA is a chromatin-associated factor that may bind to DNA in a manner dependent on other factors. We provide evidence that AtrR and FfmA interact in the cell and can influence one another’s expression. This interaction of AtrR and FfmA is required for normal azole resistance in *A. fumigatus*.

## Introduction

Resistance to the azole class of antifungal drugs is a serious clinical issue in the major filamentous fungal pathogen *Aspergillus fumigatus* (reviewed in (Wiederhold and Verweij 2020)). A likely contributor to the increased incidence of *A. fumigatus* infections associated with azole resistant isolates comes from the usage of large amounts of agricultural azole drugs that share a common target, the lanosterol α14 demethylase enzyme, with their clinical counterparts (recently reviewed in (Bastos *et al.* 2021)). This led to increased appearance of azole resistant isolates and ultimately the characterization of the genetic basis driving this phenotype (Howard *et al.* 2009; Camps *et al.* 2012; van der Linden *et al.* 2013).

One of the first and best characterized resistance alleles was a compound mutation in the *cyp51A* gene encoding the target enzyme (Verweij *et al.* 2007; Snelders *et al.* 2008). The two linked changes in *cyp51A* consisted of a nonsynonymous substitution of a leucine at position 98 with a histidine coupled with the duplication of 34 bp region in the promoter (TR34). Each of these changes were capable of decreasing the susceptibility of the fungus to azole drugs and their combination led to an enhanced level of resistance (Mellado *et al.* 2007). The TR34 repeat in the *cyp51A* promoter has been found to cause an increased level of gene transcription of this gene (Mellado *et al.* 2007). The level of *cyp51A* transcription is a key determinant of azole resistance and is determined in part by the cooperative function of the transcriptional activator proteins SrbA and AtrR (Kuhbacher *et al.* 2022; Paul *et al.* 2022c). SrbA was discovered on the basis of its similarity to the sterol response element-binding protein from mammalian cells and is a key regulator of ergosterol biosynthetic gene expression (Willger *et al.* 2008). AtrR was first described as an activator of ABC transporter-encoding gene expression (Hagiwara *et al.* 2017) and then found to be required for activation of *cyp51A* gene transcription through binding to a DNA sequence element termed the AtrR response element (ATRE) present in the 34 bp repeat (Paul *et al.* 2019).

A screen of the library of *A. fumigatus* transcription factor gene disruptions led to the identification of a C2H2 protein called FfmA as a regulator of azole resistance, in part due to its effect on transcription of the *abcG1* gene (Furukawa *et al.* 2020) (Paul *et al.* 2022a). The *abcG1* locus encodes an ABC transporter protein that is required for normal azole resistance and also a key transcriptional target of the AtrR protein (Paul *et al.* 2019). An *ffmAΔ* strain exhibited very low expression of *abcG1*, a phenotype that could be suppressed when AtrR was expressed from the strong *hspA* promoter (Paul *et al.* 2022a). The presence of this *hspA-atrR* gene fusion also rescued the slow growth phenotype caused by loss of FfmA protein further illustrating the strong genetic interaction between FfmA and AtrR.

Here we used a doxycycyline-repressible form of the *ffmA* gene to determine the transcriptome that is impacted by loss of this protein. We also carry out chromatin immunoprecipitation coupled with high throughput sequencing (ChIP-seq) to identify the direct binding sites for FfmA in the *A. fumigatus* genome. Loss of FfmA triggered changes in roughly 2000 genes, consistent with this factor controlling expression of a large portion of the *A. fumigatus* genome. We found that while 500 of these genes also were bound by FfmA in vivo, 300 of these also contained AtrR binding sites; again consistent with an interaction between these two transcriptional regulators. Expression data indicate that FfmA and AtrR regulate one another’s production and these two factors can associate in the cell as determined by co-immunoprecipitation analysis. Surprisingly, the distribution of the ATRE and FfmA binding regions across genes was found to be very different. Together our data indicate extensive interactions between FfmA and AtrR being required for the normal transcriptional program in *A. fumigatus* but these factors control gene expression in different ways.

## Materials And Methods

### *S*trains, growth conditions, and transformation

All strains used in this study were derived from AfS35 strain (*akuAΔ::loxP*). The Dox-Off-ffmA_Ble (SPF182) and the Dox-Off-FLAG-*ffmA*_Ble (SPF185) used in this study has been described in (Paul *et al.* 2022a). *A. fumigatus* strains were typically grown at 37°C in rich medium (Sabouraud dextrose; 0.5% tryptone, 0.5% peptone, 2% dextrose [pH 5.6 ± 0.2]). Selection of transformants used minimal medium (MM; 1% glucose, nitrate salts, trace elements, 2% agar [pH 6.5]); trace elements, vitamins, and nitrate salts are as described in the appendix of reference (Kafer 1977), supplemented with 1% sorbitol and either 50 mg/liter phleomycin (after adjusting the pH to 7) or 200 mg/liter hygromycin Gold (both InvivoGen). For solid medium, 1.5% agar was added. Dox off promoter shut off experiments were performed by adding 25 mg/liter doxycycline (BD Biosciences). Transformation and generation of *hasB* deletion mutants was done using in vitro-assembled cas9-guide RNA-ribonucleoproteins coupled with 50 bp microhomology repair templates (Al Abdallah *et al.* 2017). For generation of *hasB* deletion mutants, 2 CRISPR RNAs (5’-CACGAGAGTAGGATGCATGG corresponding to 5’ end of the gene and 5’-TGATGTAGGGTAGATACCAG corresponding to 3’ end of the gene) were used to replace *hasB* with the hygromycin resistance marker cassette amplified from the plasmid pSP62 (Paul *et al.* 2019)) using ultramer grade oligonucleotides from IDT harboring 50 bp homology to the break junctions of *hasB* gene. Transformants were genotypically confirmed by diagnostic PCR of the novel upstream and downstream junction formed upon targeted integration. At least 3 independent targeted transformants were phenotyped for all *hasBΔ*::hyg mutants, of which two representative strains are depicted in the data presented.

### Spot assay

Fresh spores of *A. fumigatus* were suspended in 1x phosphate-buffered saline (PBS) supplemented with 0.01% Tween 20 (1x PBST). The spore suspension was counted using a hemocytometer to determine the spore concentration. Spores were then appropriately diluted in 1x PBST. ∼100 spores (in 4 ml) were spotted on minimal medium in the presence or absence of 25 mg/ml doxycycline. The plates were incubated at 37°C and scanned after 3 days.

### Immunoprecipitation and Western blotting

Western blotting and immunoprecipitation was done as described in (Paul *et al.* 2022b) with the following modifications. For Western blotting, the AtrR polyclonal antibody used here has been described previously (Paul *et al.* 2019) and used at a 1:500 dilution. For immunoprecipitation, anti-FLAG M2 monoclonal antibody (Sigma-F1804) was used at a 1:1000 dilution.

### RNA-seq

RNA was prepared as described in reference (Paul *et al.* 2017) with the exception that cell lysates for RNA preparation were done as described above for generating samples for qRT-PCR. RNA samples were quantified using fluorimetry and RNA quality was assessed using the Agilent BioAnalyzer 2100. Sequencing libraries were generated using the Illumina TruSeq Stranded mRNA sample preparation kit (Illumina, Inc., San Diego, CA) and following Illumina’s sample preparation guide, started with 500 ng of input total RNA. The molar concentrations of the indexed libraries were measured using the 2100 Bioanalyzer (Agilent Technologies, Santa Clara, CA) with a High Sensitivity chip, and the libraries were combined equally into one pool. The molar concentration of the pool was measured using the KAPA Illumina Library Quantification Kit (KAPA Biosystems, Wilmington, MA) and the pool was sequenced on one lane of the Illumina HiSeq 4000 sequencer with a 75 bp Paired-End SBS chemistry (Illumina).

### ChIP-seq

Chromatin immunoprecipitation was done as described before (Hagiwara *et al.* 2017) with the following modifications. 25 μl was reserved as input control (IC) fraction for reverse crosslinking to verify sonication and as control for single gene ChIP and qPCR. The sheared chromatin was incubated with either anti-FLAG M2 monoclonal antibody (Sigma-F1804) at 1:250 dilution or with anti-FfmA polyclonal antibody (referenced above) at a dilution of 1:50 overnight (16 h) on a nutator at 4^0^C. This sample was further incubated with 50 μL of washed dynabeads (Life Technologies) conjugated to either protein G (when using anti-FLAG) or protein A (when using anti-FfmA) for another 8 h. Purified ChIP-ed DNA was then subjected to library preparation as outlined in (Paul *et al.* 2014) with the following modifications. The purified ChIP-ed DNA was concentrated to 10 μl using a speed-vac concentrator and the libraries generated using the Rubicon ThruPLEX DNAseq kit using 10 μl as input volume.

### Data availability statement

Strains, antibodies and plasmids are available upon request. ChIP- and RNA-seq data are available at GEO with the accession number: GSE229913.

## Results

### Relationship between *S. cerevisiae* Mot3 and FfmA

Inspection of the Panther protein sequence database (Thomas *et al.* 2022) provided an indication that the overall structure of FfmA was similar to that of the Mot3 transcriptional regulator from *S. cerevisiae*. The two C2H2 domains of FfmA are located in the center of the factor (Figure 1A). Alignment with Mot3 indicated these proteins also exhibited sequence conservation with most of this centered around the C2H2 domains (Figure 1B). Importantly, while both these factors contain C2H2 domains, there are many substitutions and spacing changes between these domains. Note that the extensive poly-asparagine N-terminus of Mot3 is not retained in FfmA while the C-terminus of FfmA extends beyond that of Mot3. Previous work has implicated Mot3 (<u>M</u>odulator <u>o</u>f transcription) as a sequence-specific DNA-binding protein involved in regulation of a wide range of different genes including ones involved in mating pheromone response (Grishin *et al.* 1998), ergosterol gene expression (Hongay *et al.* 2002), anaerobic gene regulation (Kastaniotis *et al.* 2000), and control of transposon gene expression (Madison *et al.* 1998). We present data supporting the view that while *A. fumigatus* FfmA also acts as a regulator of gene transcription impacting a wide range of genes as does Mot3, the mechanisms used by these two structurally related proteins are not the same.

**Figure 1.**
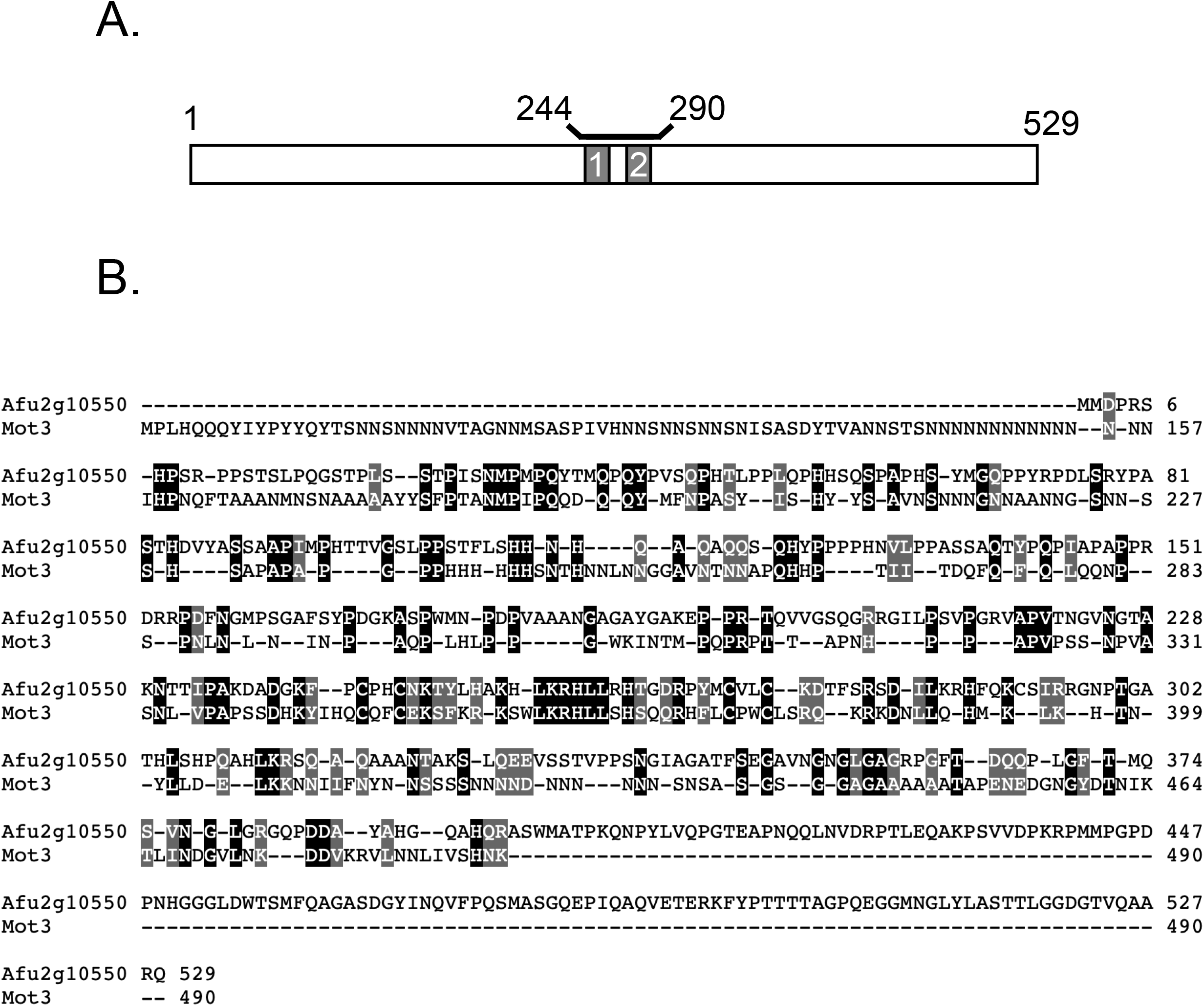
Structure of FfmA. A. A cartoon depiction of the structure of FfmA is shown with the two C2H2 domains indicated as boxes 1 and 2. These two domains are located between amino acid residues 244 and 290. B. Sequence alignment of FfmA with S. cerevisiae Mot3. Identical residues are boxed in black, conservative substitutions are boxed in gray and gaps are indicated by the dashes.

### Chemical repression of *ffmA* gene expression

Previous characterizations of loss of *ffmA* function demonstrated that chronic loss of function mutant strains lacking this gene had severe growth defects (Liu *et al.* 2021; Paul *et al.* 2022a). To limit indirect changes caused by general defects in growth, we turned to the use of a doxycycline-repressible allele of *ffmA* that we have evaluated in earlier work (Paul *et al.* 2022a). The allele was produced by inserting a doxycycline-regulated promoter into the *ffmA* locus such that it was downstream of the normal *ffmA* regulatory region and now responsible for production of *ffmA* mRNA and protein. To examine the rate of loss of FfmA protein from doxycycline-treated cells, parallel cultures of the Dox off-*ffmA* strain were grown in the absence of doxycycline for 12 hours and then treated with 25 mg/l of this drug over an 18 hour time course. Total protein extracts were prepared and analyzed by western blotting (Figure 2A)

**Figure 2.**
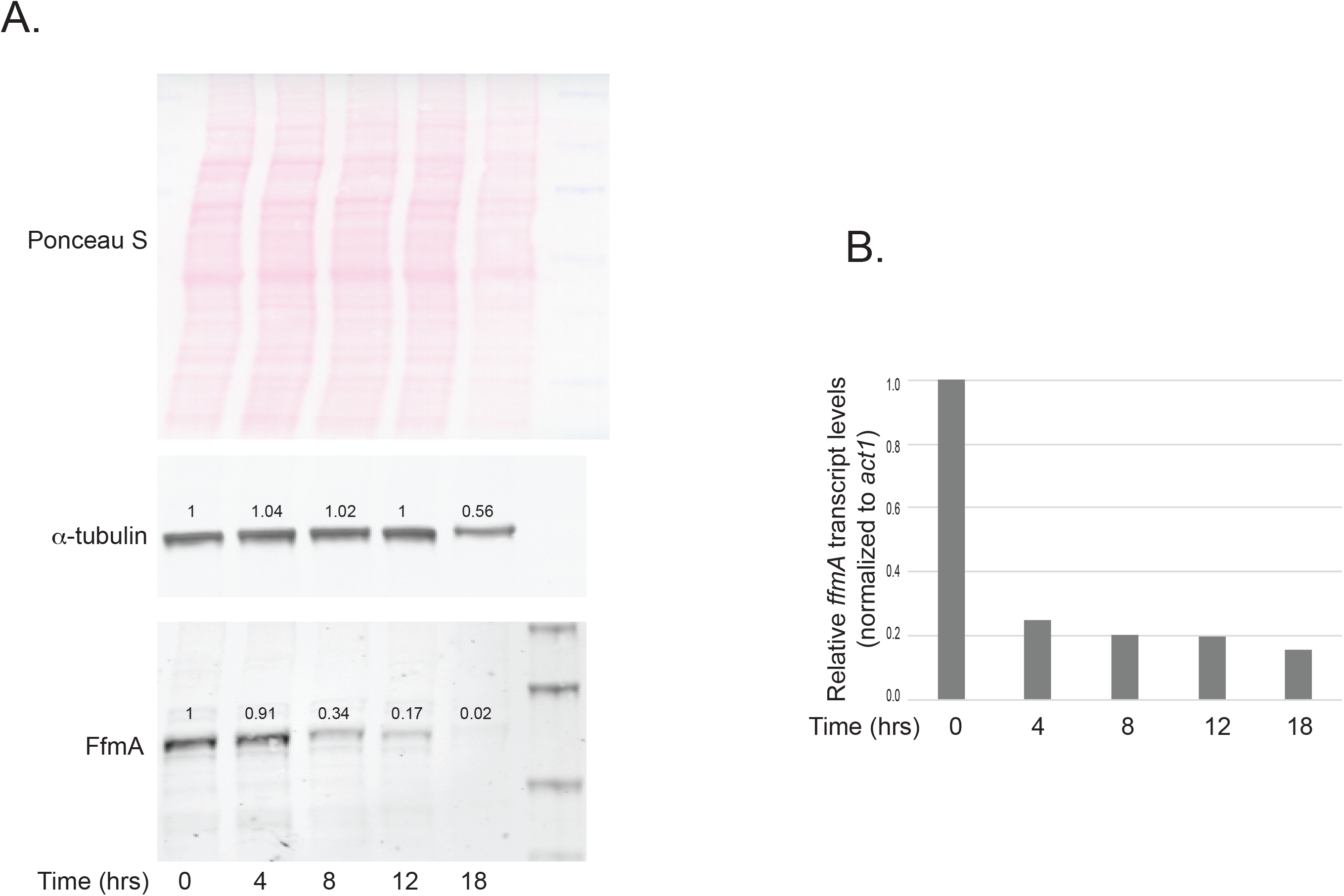
Doxycycline-depletion of doxycycline-regulated *ffmA* transcription. A. A doxycycline-repressible (doxycycline off: DO) promoter-driven *ffmA* gene (DO-*ffmA*)- containing strain was grown overnight in the absence of doxycycline. A sample of this culture was processed to measure FfmA expression in the absence of doxycycline (0 hours) and then doxycycline was added for the numbers of hours indicated. Cultures were processed at each indicated time point and whole cell protein extracts prepared to determine the effect of doxycycline addition on expression of FfmA protein by western blotting with anti-FfmA. Tubulin was also analyzed as a loading control along with Ponceau S staining of the membranes after protein transfer. The numbers refer to relative expression normalized to the 0 hour time point. B. Samples from the strains grown as in A were processed for RNA and levels of either *ffmA* or *act1 mRNA* were determined by qPCR.

Levels of FfmA protein were lowered to roughly 34% of normal after an 8 hour incubation in the presence of doxycycline. This reduction continued at 12 and 18 hours to 17 or 2% of normal expression, respectively. We used two different loading controls to determine if any generalized reduction in protein levels could be detected. Levels of tubulin protein were stable over the entire time course until 18 hours after the addition of doxycycline. At this point, tubulin levels were seen to drop to 60% of the untreated culture. Additionally, total protein measured by Ponceau S staining of the membrane prior to transfer indicated a reduction in overall protein levels. We interpreted these data as consistent with a generalized defect in protein synthesis that occurred somewhere after 12 hours of repression of *ffmA* transcription.

We also assessed the effect of doxycycline addition on the DO-*ffmA* allele by preparing RNA from each of the time points above and measuring *ffmA* mRNA levels using a qRT-PCR assay (Figure 2B). Levels of *ffmA* transcript dropped after 4 hours of growth in the presence of 25 mg/l doxycycline and stayed low throughout the time course. These experiments demonstrate that the DO-*ffmA* allele could be used to more acutely deplete both FfmA protein and RNA from cells prior to analyzing the biological consequences of this loss.

### Acute depletion of FfmA caused large changes in the *A. fumigatus* transcriptome

Parallel cultures of the DO-*ffmA* strain were grown for 12 hours in the absence of doxycycline and then 25 mg/l of this drug was added to one culture. Both were grown for an additional 8 hours and total RNA prepared. We also grew wild-type cells as a control for normal gene expression. RNA from these 3 strains was then used in an RNA-high-throughput Sequencing (RNA-seq) experiment to profile the transcriptomes produced (Figure 3A). After calculation of the FPKM value, we used DEseq2 to produce a ratio of RNA expression values in each experimental condition relative to that seen in the wild-type strain. We limited our general conclusions to genes that were at least log2=1 changed in expression with a P<0.05.

**Figure 3.**
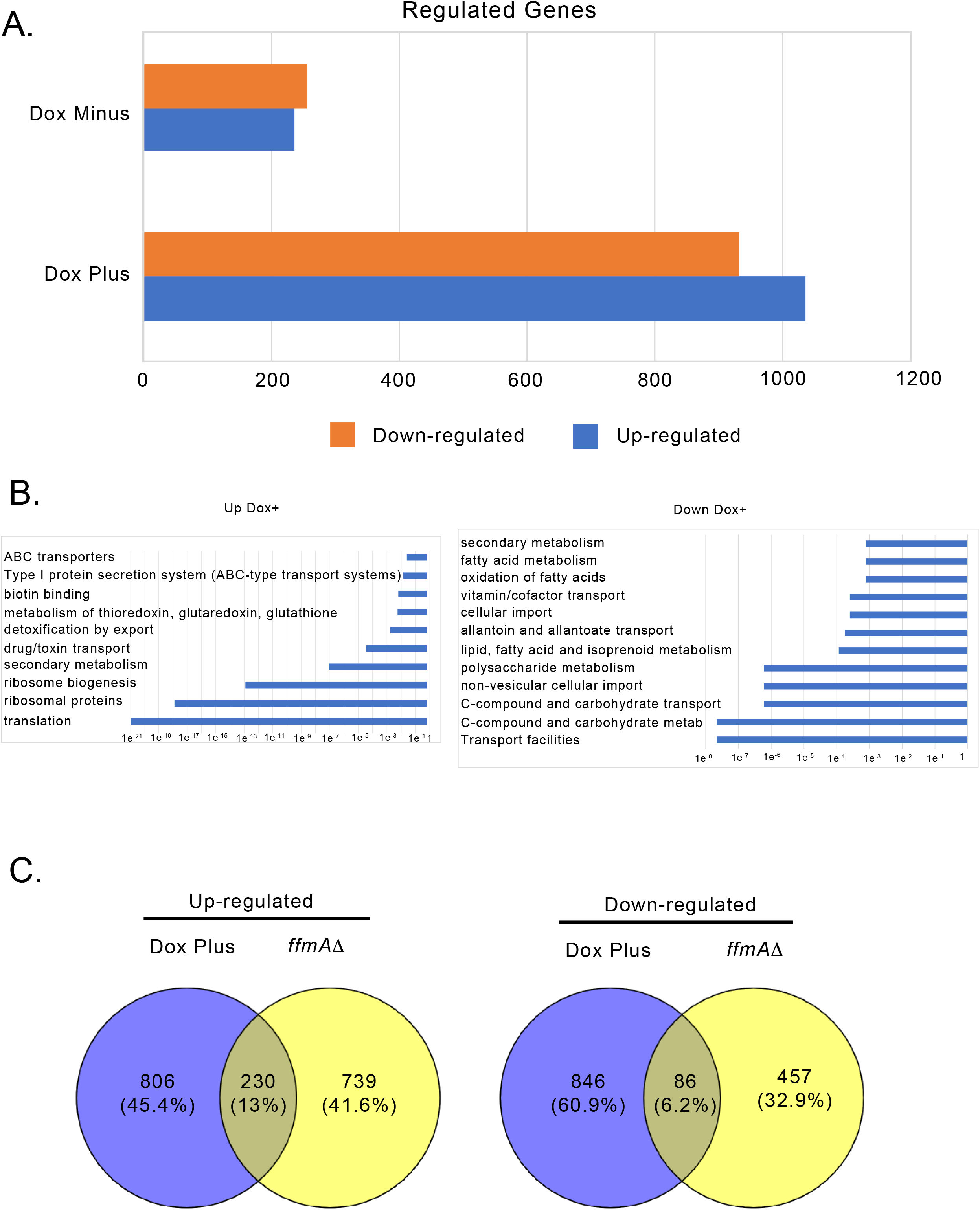
RNA-seq analysis of doxycycline-regulated *ffmA* gene expression. A. The numbers of genes significantly up- or down-regulated by at least two-fold in the presence of the doxycycline-repressed (dox off: DO) DO-*ffmA* allele are quantitated under conditions in which FfmA expression is high (Dox Minus) or after doxycycline addition to lower FfmA levels (Dox Plus). B. Significant categories of genes either induced (Up Dox+) or repressed (Down Dox+) by FfmA in the presence of doxycycline. C. Number of genes that are commonly regulated between the doxycycline-inhibited DO-*ffmA* strain and earlier experiments using a *ffmAΔ* allele (Paul *et al.* 2022a).

Growth of the DO-*ffmA* strain in the absence of doxycycline led to similar numbers of genes that were either up-regulated (236) or down-regulated (259). Since growth of the DO-*ffmA* strain under this condition was closer to that of the wild-type than in the case of the *ffmAΔ* null strain, we anticipated that the transcriptome would also be less disturbed. Using FungiFun2 (Priebe *et al.* 2015) to perform FunCat (Ruepp *et al.* 2004) classification, we examined the function of the genes seen to be co-regulated under these conditions and were surprised to find that the genes that were increased in expression compared to the wild-type strain did not show any significant enrichment of any functional class. The presence of the DO-*ffmA* allele led to a log2 of 1.6 for the *ffmA* mRNA itself, indicating that this construct elicited overproduction of *ffmA* (Supplementary table 1). Conversely, the genes that were reduced in expression compared to wild-type did show significant enrichment and were primarily concerned with metabolism and cell transport (Supplementary Table 2). We did notice a modest growth defect in this strain in the absence of doxycycline but its growth was dramatically improved from that of the *ffmAΔ* null (Liu *et al.* 2021; Paul *et al.* 2022a).

Strikingly, the addition of doxycycline to this DO-*ffmA*-containing strain led to a much larger change in gene expression, consistent with the major reduction in growth rate caused by acute repression of *ffmA*. 1036 genes were induced upon doxycycline addition while a similar number (932) were repressed (Supplementary table 3). Together, this represents nearly 20% of the *A. fumigatus* genome, supporting a global effect caused by depletion of FfmA upon doxycycline repression.

The three major classes of genes that were induced upon doxycycline repression of *ffmA* expression were associated with translation and ribosomes, secondary metabolism and transport across membranes (Figure 3B; Supplementary table 4). Two small groups of genes encoding proteins involved in production of antioxidants as well as metabolic enzymes that use biotin as a cofactor were also induced. It was surprising that loss of normal FfmA expression had such a striking induction of components of the translational apparatus. Typically, cells repress biosynthesis of the translation machinery when undergoing stress (reviewed in (Holcik and Sonenberg 2005)). We will discuss a possible explanation for this transcriptional response below.

Secondary metabolism genes and transport proteins were also strongly induced when FfmA levels dropped. Our original interest in FfmA was prompted by its effect on expression of the ATP-binding cassette transporter AbcG1 (Paul *et al.* 2022a) and these RNA-seq data indicated that at least 8 different ABC transporter genes were activated when FfmA was repressed. When all membrane proteins were considered, nearly 40 genes were induced upon loss of normal FfmA expression. In the large number (145) of secondary metabolic genes that were induced in doxycycline-treated DO-ffmA cells, 25% of these genes were also membrane proteins. These results suggest that one important role of FfmA is to regulate expression of membrane transporter proteins, potentially in response to changes in accumulation of secondary metabolic intermediates.

While a large number of genes were also repressed on the addition of doxycycline to the DO-*ffmA* strain, the maximum significance of these was less than for those genes that were induced (e^-8^ compared to e^-21^, respectively). These repressed genes could be broadly separated into loci involved in either transport or metabolism (Supplementary table 4). These repressed transporters were classed as importers of nutrients rather than the exporters seen to be induced above. As with the induction of the translational components, this was surprising as the metabolic stress being triggered by loss of FfmA might be expected to trigger an increased need for nutrient uptake. Similarly, loss of metabolic gene expression was extensive and could be the result of FfmA depletion stress causing metabolic rewiring.

We also compared these acute changes above with the chronic changes in gene expression seen previously in RNA-seq comparison of the transcriptomes of wild-type and *ffmAΔ* null strains (Liu *et al.* 2021). There was limited overlap between these gene sets: 13% of genes were shared in terms of loci induced in *ffmAΔ* null cells compared to the doxycycline-treated DO-*ffmA* strain while 6% were shared in the repressed gene sets (Figure 3C). The top two classes of shared induced genes were involved in secondary metabolism and drug/toxin export (Supplementary table 5). A noticeable absence from these shared genes were those encoding components of the translational machinery. During the chronic lack of FfmA, these genes are no longer overproduced as they are upon the acute absence of this factor. Shared repressed genes are largely membrane proteins, often thought to be involved in nutrient uptake (Supplementary table 5).

Genes that were differentially regulated in either the chronic loss of FfmA or during doxycycline repression were also found. Genes induced in the *ffmAΔ* null strain but repressed in the doxycycline-treated DO-*ffmA* strain include a large collection of membrane importers and also a group of metabolic enzymes involved primarily in carbon metabolism (Supplementary table 5). Possibly the long-term adjustment to loss of FfmA leads to a change in fundamental metabolic processes as has been suggested earlier (Liu *et al.* 2021).

Finally, a smaller collection of genes that were elevated in doxycycline-repressed DO-*ffmA* cells but repressed in the *ffmAΔ* background was also identified. These genes only included 4 membrane transporters, all of which were involved in secondary metabolism, while the rest were involved in direct metabolic events including urea metabolism or were classified as defense-related proteins (Supplementary table 5). Interestingly, *abcG1* fell into this category as its log2 change was 0.74 in doxycycline-treated DO-*ffmA* and -0.5 in *ffmAΔ* cells. The magnitude of these changes was below our cutoff of log2=1 but fully consistent with our previous data (Paul *et al.* 2022a). This pattern of gene expression suggests that these factors are involved in the short-term adjustment to loss of FfmA but might be eventually repressed when its loss is longer term.

Taken together, these transcriptomic data indicate that FfmA controls a large suite of genes and seems to act both positively and negatively on their expression. No simple, overarching theme emerged from these FfmA responsive genes. In our previous studies on AtrR, we found that a minority of genes that were transcriptionally responsive to this upstream binding factor actually contained AtrR response elements (ATREs), the recognition site for this factor (Paul *et al.* 2019). We wanted to determine what fraction of these FfmA-responsive genes were direct target genes of this factor through the use of chromatin immunoprecipitation-high-throughput DNA sequencing (ChIP-seq).

### Genomic binding sites of FfmA

We performed ChIP-seq analysis of FfmA DNA-binding using two different immunological reagents to recover FfmA-enriched chromatin. We used a previously developed anti-FfmA antiserum to detect native protein as well as a FLAG-tagged FfmA protein driven from the doxycycline-repressible promoter. Identical ChIP-seq experiments were performed using these antisera and either a wild-type strain or the DO-FLAG-ffmA strain. After mapping the peaks of FfmA binding seen in these two different experiments, we collected the overlapping peaks to focus on those most likely to bind FfmA in vivo (Supplementary table 6).

616 different peaks were identified in this analysis, representing 530 unique genes. No significant genes were found to be enriched via FunCat analysis (Data not shown). Compared to our previous analysis of the ChIP-seq profile of AtrR, we noticed many more peaks for FfmA inside or downstream of the coding sequence (∼40%) compared to those for AtrR (∼22%) (Paul *et al.* 2019). These data represented the first indication that FfmA was not likely to act as a typical upstream activation sequence-binding protein as we had found for AtrR. Several different FfmA target genes are shown to compare the binding of AtrR to that seen for FfmA, along with the RNA-seq data from wild-type and DO-ffmA +/- doxycycline (Figure 4A).

**Figure 4.**
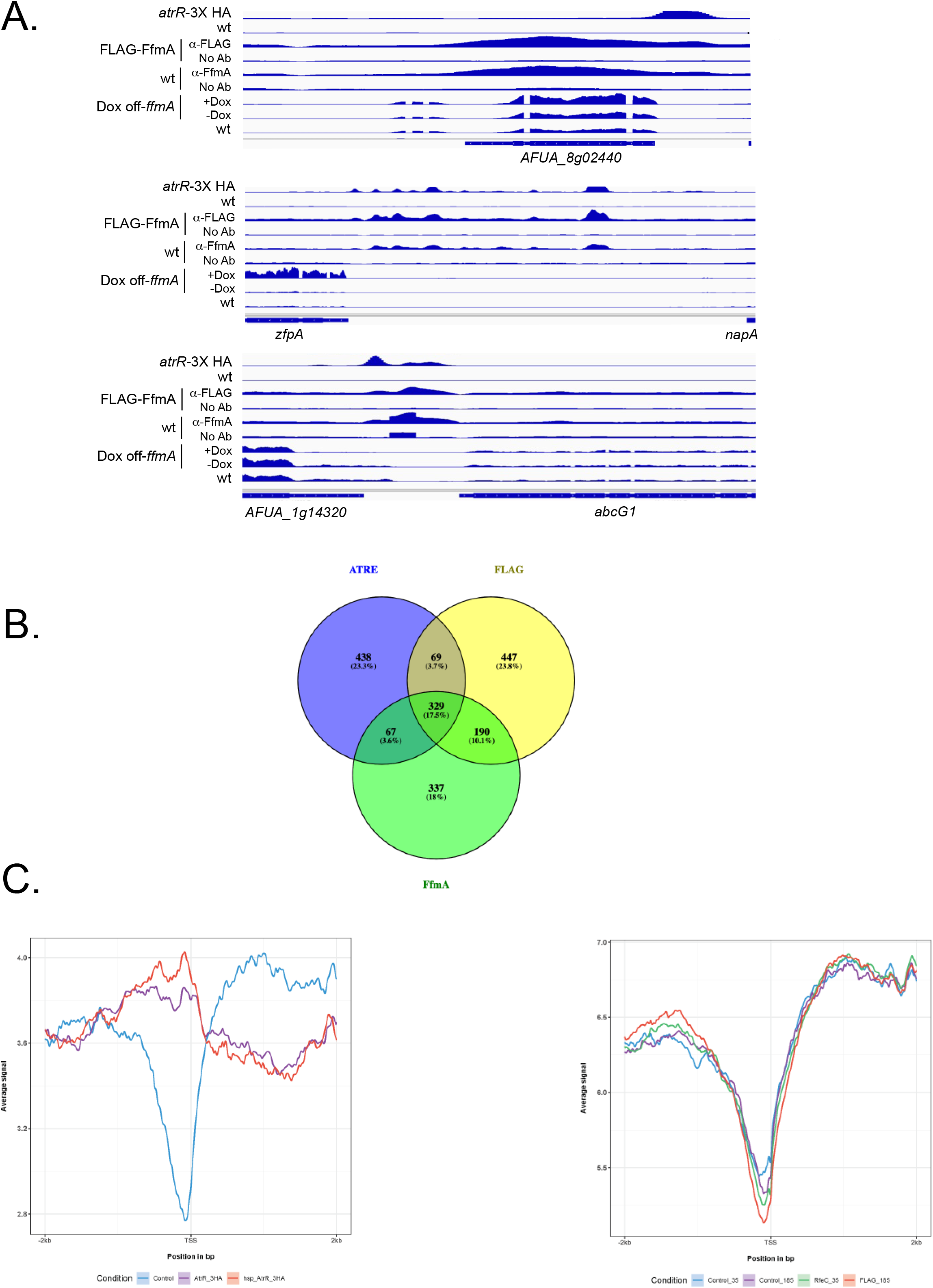
Chromatin immunoprecipitation-high throughput sequencing (ChIP-seq) analysis of FfmA. A. Integrative Genetics Viewer (IGV) plots of ChIP-seq data for AtrR, FfmA and RNA-seq data for FfmA are shown. ChIP-seq data of AtrR was taken from (Paul *et al.* 2019) and HA antibody was used to recover chromatin from isogenic cells containing or lacking an *atrR*-3X HA fusion gene. FfmA ChIP-seq data were generated using anti-FLAG antibody to isolate chromatin from a strain containing or lacking a FLAG-ffmA fusion gene (FLAG-ffmA). Additionally, chromatin was isolated from a wild-type strain using either an anti-FfmA polyclonal antibody or with no primary antibody (No Ab). RNA-seq data are shown for RNA isolated from a wild-type (wt) strain or from the DO-ffmA strain grown in the absence (-Dox) or presence (+Dox) of 10 mg/l doxycycline. B. Overlap in genes with ATREs and exhibiting ChIP-seq peaks for immunoprecipitated chromatin performed with either anti-FLAG or anti-FfmA as detailed above. C. RECOUP plots of ChIP-seq data from either AtrR (left hand panel) or FfmA (right hand panel). These plots were generated using ChIP-seq data for all genes from - 2 kb upstream and downstream of the transcription start site (TSS) and represent the average occupancy for either AtrR or FfmA across all genes in *A. fumigatus* larger than 1 kb (to aid in peak separation between 5’ and 3’ ends of genes). These data correspond to ChIP-seq analysis of strains expressing two different forms of *atrR*-3X HA (either driven by the wild-type promoter or the strong *hspA* promoter) as well as a strain lacking any HA tag. The two different means of detecting FfmA-bound chromatin described above as well as their controls are plotted on the right. Note that AtrR shows the expected enrichment for a factor that binds to the 5’ end of genes while FfmA shows no such enrichment and is even more likely to bind to the region downstream of the TSS.

The top panel illustrated data from the *AFUA_8g02440* (*erg25A*) gene in which AtrR binds upstream of the transcription start site but the shared FfmA peaks are located within the transcription unit. The transcription factor-encoding *zfpA* gene contains a number of AtrR peaks in its promoter region and these are closely paralleled by the appearance of FfmA binding density. Note the strong induction of *zfpA* transcription upon addition to doxycycline to the DO-*ffmA* gene. Finally, the *abcG1* gene is shown as the effect of FfmA loss of this factor caused a reduction in AbcG1 expression by western blot analysis (Paul *et al.* 2022a). As was seen for *zfpA*, FfmA peaks are upstream of the *abcG1* transcription. For both *zfpA* and *abcG1*, the regions bound by FfmA and AtrR overlap, a configuration that would be expected if both these factors were upstream transcriptional regulators. Note that this is not the case for *erg25A* that provides an illustration of the fundamental differences between FfmA and AtrR function. Our analyses of the total spectrum of FfmA and AtrR binding sites indicates that the majority of joint FfmA and AtrR target genes are represented by the *erg25A* locus (see below).

We also compared the genes that were either significantly induced or repressed in DO-ffmA cells treated with doxycycline with the 530 genes showing binding of FfmA using the two different immunological reagents to perform ChIP-seq. Only around 20% of the 530 genes exhibited altered gene expression under these conditions and of these, only 6 genes were seen to be significantly enriched in the list of genes induced in the presence of doxycycline in DO-ffmA cells. These genes were all membrane transporter proteins and an amine oxidase protein (Supplementary table 7). As we found for AtrR, the majority of genes altered by changes in FfmA dosage occur via indirect effects on gene expression.

To determine the total extent of overlap between FfmA and AtrR binding sites in the *A. fumigatus* genome, we compared the genes that were detected in the two different ChIP-seq experiments for FfmA with our previous data for AtrR (Figure 4B). 329 genes were detected that were bound by FfmA and AtrR. Analysis by FunCat did not show any shared functions of these genes but GO term analysis in FungiDB indicated that FfmA and AtrR bound to 58 different genes classified as being involved in regulation of nucleic acid-templated transcription (Supplementary table 8). Additionally, another highly significant enriched GO term category was asexual sporulation (Supplementary table 9). There were 14 genes in this category and at least 8 of these were also transcription factors. Clearly, FfmA and AtrR share many target genes that encode transcriptional regulatory proteins.

Given the degree of shared target genes between FfmA and AtrR, we wanted to compare the genomic coverage from our ChIP-seq assays of FfmA and AtrR to determine if similar binding profiles occurred for these factors on their target genes. We used a software package called Recoup (Moulos 2021) to generate an average coverage profile of genome-wide binding across the *A. fumigatus* genome for both FfmA and AtrR target genes. These profiles were oriented relative to the annotated transcription start site (TSS) and restricted to genes that were at least 1 kb in length to enhance separation between promoter regions and the subsequent intergenic region. These plots are shown in Figure 4C.

AtrR ChIP-seq coverage showed the profile expected of an upstream binding transcriptional regulator with its coverage peaking from -1 kb 5’ of TSS and decreasing to a near minimum close to the transcription start site (Figure 4C, left panel). Strikingly, FfmA showed a very different profile with maximum coverage downstream of the TSS and the lowest coverage from the TSS to 1 kb upstream. FfmA binding seemed to be distributed fairly evenly between regions upstream and downstream of the TSS, quite differently from AtrR. The difference in the global coverages seen between all AtrR target sites compared to those of FfmA target genes provides the clearest indication that these two factors do not regulate transcription in an identical manner.

While the mode of action of FfmA and AtrR may be different based on their genome coverage, there is evidence for many shared target genes as illustrated by the ChIP-seq data above. To determine if these factors might interact at other levels, we analyzed their expression patterns and protein:protein interaction.

### Interaction between FfmA and AtrR proteins

To assess if changing the dosage of either *ffmA* or *atrR* might influence the expression of the other factor, we used western blot analysis to compare FfmA expression isogenic wild-type and *atrRΔ* strains. Whole cell protein extracts were prepared from these two strains and then analyzed by western blotting using polyclonal α-AtrR antiserum (Figure 5A).

**Figure 5.**
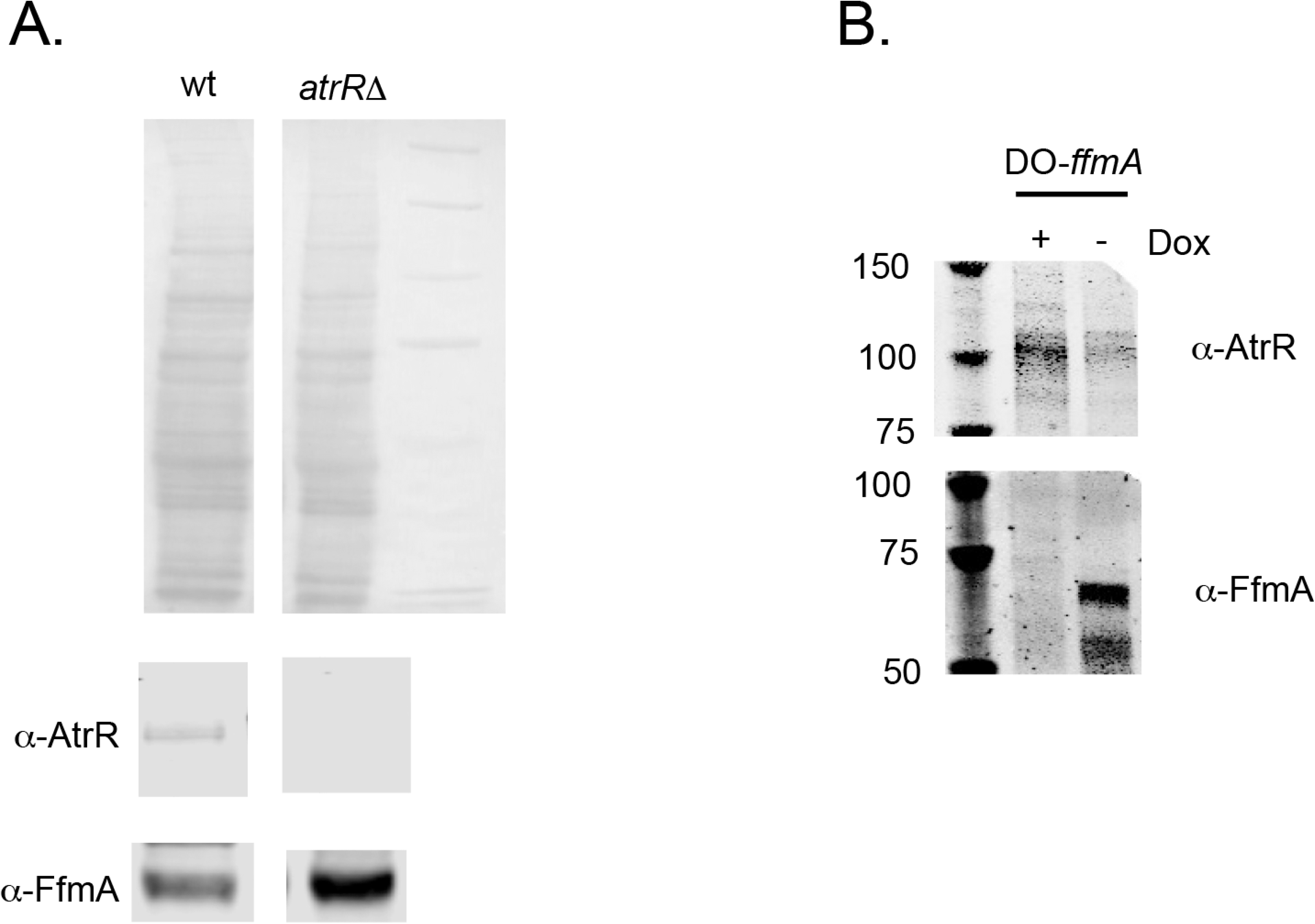
Interaction at the level of expression between AtrR and FfmA. A. Isogenic wild-type and *atrRΔ* strains were grown and whole cell protein extracts prepared. Equal amounts of these extracts were analyzed by SDS-PAGE using either anti-AtrR or anti-FfmA antisera. B. The DO-*ffmA*-containing strain was grown either in the presence (+) or absence (-) of doxycycline for 12 hours. Whole cell protein extracts were analyzed as described above using the same antibodies. Molecular mass standards are indicated in kD on the left.

Loss of AtrR triggered an increase in the steady-state level of FfmA protein, indicating that the accumulation of FfmA is sensitive to the levels of AtrR. Conversely, we used the DO-*ffmA* strain to acutely deplete FfmA production and assessed AtrR levels both in the presence and absence of doxycycline (Figure 5B).

Depletion of FfmA from the DO-*ffmA* strain upon doxycycline addition led to the induction of AtrR. Together, these data indicate that expression of AtrR and FfmA are mutually antagonistic and support the notion that a regulatory interaction exists linking these two factors. This regulatory interaction, coupled with the high degree of shared ChIP-seq peaks between AtrR and FfmA, led us to test if these two factors might interact. Previous mass spectrometric experiments suggested that FfmA could be co-purified with AtrR (Paul *et al.* 2022b). We directly tested this idea using co-immunoprecipitation analysis.

The DO-FLAG-*ffmA* strain was grown overnight and whole cell protein extracts prepared under nondenaturing conditions. An aliquot of this extract was reserved as input control. The remaining extract was immunoprecipitated with either no primary antibody (No Ab) or with anti-FLAG antibody. A secondary anti-mouse antibody bound to beads was used to recover immunoprecipitates. These were washed and then resolved on SDS-PAGE. Recovered proteins were detected by western blotting (Figure 6).

**Figure 6.**
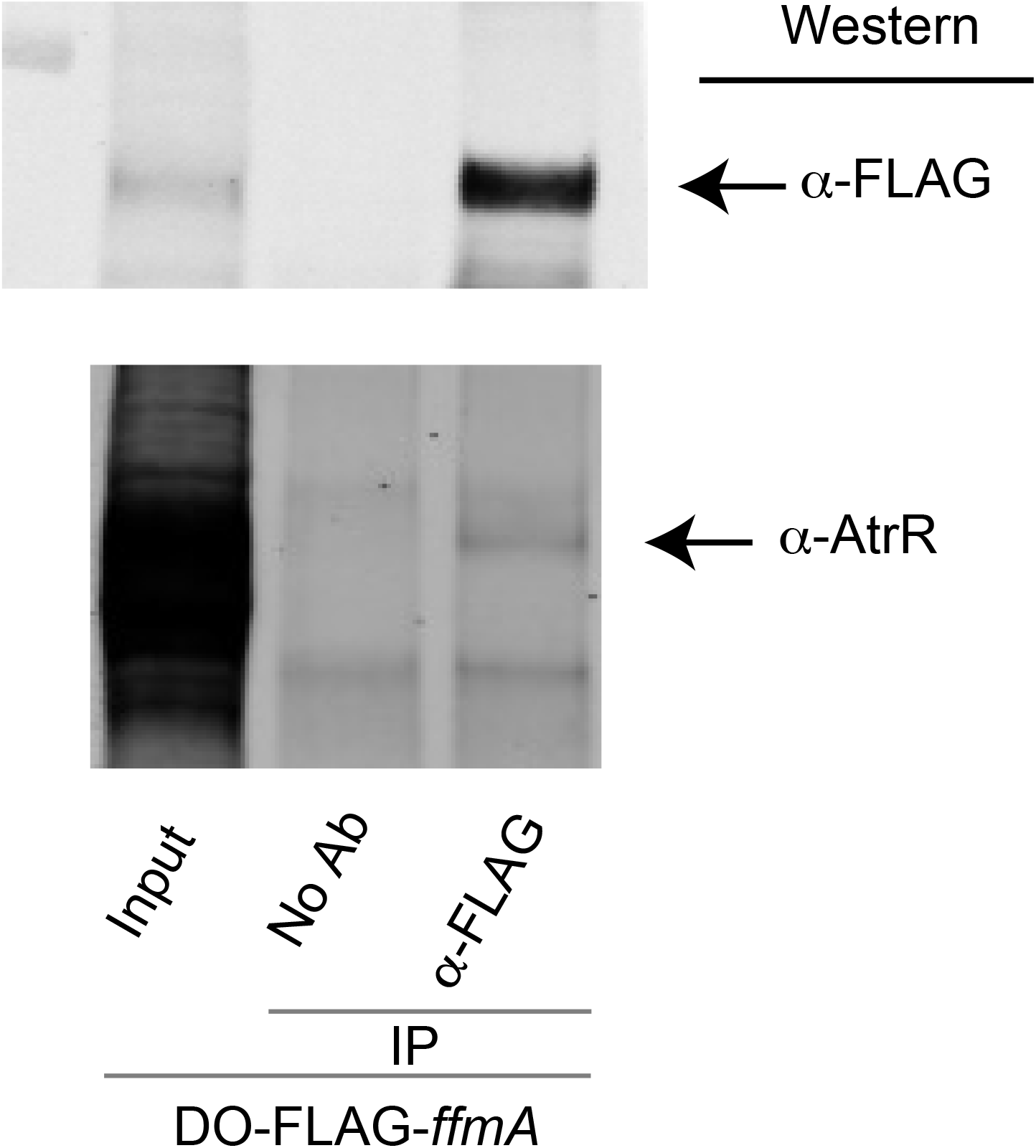
AtrR interacts with FfmA. The DO-FLAG-ffmA strain was grown and whole cell extracts prepared under nondenaturing conditions. An aliquot of this original extract was retained and used as an input control (Input). Immunoprecipitations were conducted with anti-FLAG antibody or with no primary antibody (No Ab). Immunoprecipitates were washed and then denatured. Aliquots of immunoprecipitated proteins were analyzed by western blotting using either anti-AtrR or anti-FLAG antibody.

We were able to detect AtrR in the FLAG immunoprecipitated fractions but not in an identical immunoprecipitation lacking the anti-FLAG primary antibody. These data support the idea that AtrR can interact with FfmA in the cell. Note that only a small amount of AtrR protein was recovered in the anti-FLAG immunoprecipitate of FLAG-FfmA. This supports the view that while AtrR does with FfmA, this represents a minor fraction of total AtrR. We will discuss the implications of this finding below.

### Expression of an iron chelator upon loss of FfmA impacts pigmentation phenotype

Previously, we had noticed that strains lacking FfmA accumulated an orange pigmentation during growth along with a profound growth phenotype. Other experiments demonstrated that the *has* gene cluster produces an iron chelator called hexadehydroastechrome that acts to link iron levels with secondary metabolite production (Wiemann *et al.* 2014). Overproduction of *has* genes via overexpression of the HasA transcription factor, with concomitant elevation of hexadehydroastechrome levels also caused an increase in expression of the high-affinity iron permease *ftrA*, the siderophore *sidA* and a decrease in the iron-consuming protein cytochrome C (Wiemann *et al.* 2014). Strikingly, these high levels of *has* gene expression also caused the cells to acquire an orange pigmentation (Yin *et al.* 2013). These data are consistent with high levels of hexadehydroastechrome acting to chelate internal iron and trigger the normal cellular response to iron limitation. We found that the addition of doxycycline to the DO-*ffmA* strain also led to an induction of the *hasAB* regulon (Figure 7A) and wondered if this addition of doxycycline might cause our strain to grow slowly and acquire the orange pigmentation due to elevated has gene expression as was observed when *hasA* was overproduced (Yin *et al.* 2013). To test this idea, we disrupted the *hasB* gene in our DO-*ffmA* strain. Transformants were recovered and then placed on either minimal media with or without doxycycline.

**Figure 7.**
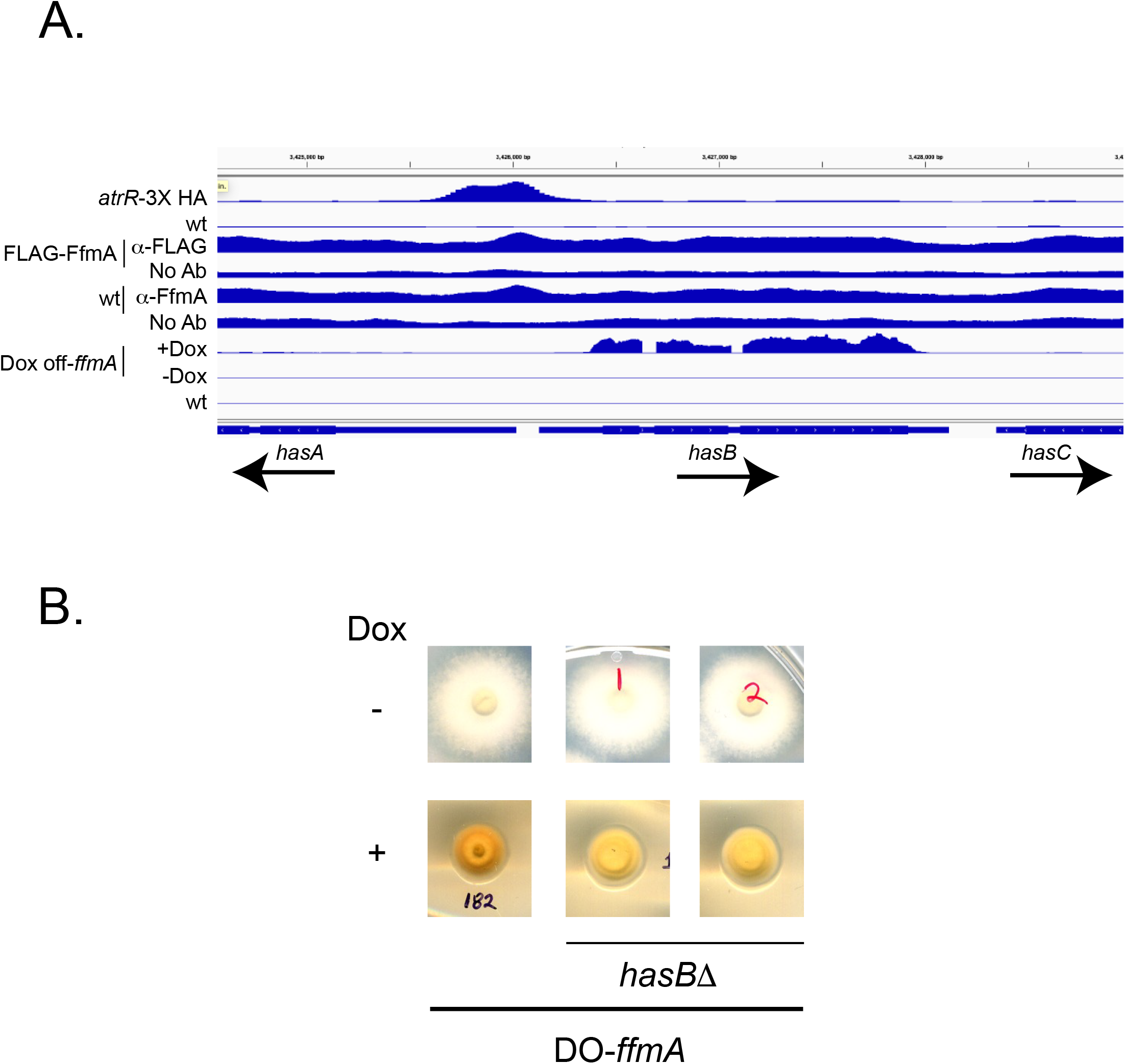
Pigmentation caused by depletion of FfmA is dependent on expression of *hasB*. A. ChIP- and RNA-seq data at the *hasA/*B locus. An IGV plot as shown in Figure 3A summarizes the location of the AtrR binding region as well as the absence of significant FfmA binding under these conditions. When FfmA is depleted by doxycycline treatment of the DO-*ffmA* strain, *hasB* transcription was seen to dramatically increase. B. Loss of *hasB* reduces the pigmentation seen in DO-*ffmA*. The *hasB* gene was disrupted with insertion of a hygromycin resistance marker and two isolates were tested for their degree of pigmentation after repression of FfmA expression by doxycycline. Note the deeper orange color of the *hasB* isolate compared to the lighter yellow color of the two *hasBΔ* strains in the presence of doxycycline.

Removal of the *hasB* gene reduced the level of orange pigmentation that was seen in DO-*ffmA* cells in which FfmA expression had been repressed with the addition of doxycycline (Figure 7B). Note the dark orange color of the DO-*ffmA* cells upon doxycycline repression compared to the isogenic *hasBΔ* derivatives. Growth of the DO-*ffmA* strain in the absence of doxycycline showed no impact on pigmentation due to the presence of the *hasB* gene. These data support the view that loss of FfmA function leads to an induction of HasB that in turn leads to increased hexadehydroastechrome chelating iron. We believe this is an indirect caused by loss of FfmA as significant binding of FfmA was not detected in the *hasAB* region (Figure 7A). This finding also illustrates the difference between FfmA and AtrR function at *hasAB* as AtrR directly binds to this promoter although FfmA does not. Even with this bound AtrR, activation of the *hasAB* regulon required depletion of FfmA from the cell.

## Discussion

### FfmA is a chromatin-associated protein that has wide-ranging effects on gene regulation

We initially hypothesized that FfmA protein would act in a fashion similar to that of AtrR (Paul *et al.* 2022a). While many sequence-specific transcriptional regulators of the C2H2 class of proteins are known (Emerson and Thomas 2009), this protein domain also can act to support protein:protein interactions (Brayer and Segal 2008) as well as to bind RNA (Lu *et al.* 2003) or methylated DNA (Prokhortchouk *et al.* 2001). We suggest FfmA associates with chromatin but does not do so in a direct sequence-specific manner based on multiple observations. The structural similarity between FfmA and Mot3 (Figure 1) suggests that while these proteins are both global transcriptional regulators, this conservation does not extend to the manner in which each associates with DNA target genes. We base this suggestion on multiple lines of evidence.

First, we were able to readily generate ChIP-seq profiles for FfmA using either a FLAG-tagged version of the protein or a rabbit polyclonal antibody directed against the native factor. The peaks from these different experiments extensively overlapped, providing confidence that these corresponded to authentic sites of FfmA enrichment in vivo. However, analysis using MEME-ChIP (data not shown) of these data sets led to the detection of an element very similar to that for the AtrR response element (ATRE). This sequence motif was also detected in a minority of FfmA ChIP-seq peaks (45/600) and was not centrally located in these peaks. This behavior is more consistent with an element peripherally associated with the binding of FfmA as an authentic binding site would be expected to be centrally located in MEME-ChIP analysis as this involves inspecting 250 bp on either side of the detected ChIP-seq peak and deriving a consensus (Ma *et al.* 2014). Typical binding sites would be expected to be centrally located by this analysis.

Secondly, efforts to detect direct DNA-binding by recombinant FfmA protein were not successful (data not shown). We were able to detect general DNA binding but no sequence specific site recognition. Conversely, this was relatively straightforward for AtrR (Paul *et al.* 2019). One possibility we cannot exclude is that we chose regions of promoters that, while corresponding to strong FfmA ChIP-seq signals, do not work well in an in vitro assay. Based on the significant number of genes (329 genes, Figure 4B) that share binding regions for both FfmA and AtrR, it is possible that direct DNA-binding of FfmA requires interaction with an additional factor.

Finally, the Recoup plotting of the ChIP-seq peaks is the most striking indication that the action of FfmA and AtrR are different. This algorithm plots the average sequence coverage across all genes by the factor of interest. If the Recoup treatment of AtrR ChIP-seq data is inspected, a clear enrichment of binding can be seen on the 5’ end of all genes (Figure 4C). Note this is quite distinct from the analysis of the FfmA ChIP-seq data as there is no apparent enrichment of FfmA binding to either side of the transcription start site. This pattern of binding is very reminiscent of that seen for the Set1 histone methyltransferase from yeast (Luciano *et al.* 2017) as well as the Gds1 protein that binds to a histone acetyltransferase (Joo and Buratowski 2022). Set1 associates with genes via interaction with RNA while Gds1 associates with a histone acetyltransferase. The basis of FfmA chromatin association is yet to be determined although it seems likely to involve AtrR, at least in part. Our co-immunoprecipitation experiments indicated that only a small fraction of FfmA was associated with AtrR, arguing that other means of chromatin binding must be present for normal FfmA function.

The chromatin-bound FfmA protein functions to control the expression of a wide range of genes. We found roughly 2000 genes that were significantly changed by at least two-fold in cells that were depleted of FfmA in an acute manner (Supplementary table 1). This large effect on genomic transcription is consistent with FfmA acting globally to impact overall gene expression rather than to control some selective group of genes. Additionally, a small fraction of these genes (∼10%) were also associated with detectable FfmA binding by ChIP-seq. Together, these data indicate that much of the effect seen upon changing dosage of FfmA is indirect. Since we found that a large number of genes encoding transcriptional regulatory proteins are bound by FfmA (Supplemental table 8), this may provide a mechanism explaining why so many genes can respond to changes in this factor with no associated direct DNA-binding.

One striking example of this could explain why so many ribosomal components are elevated upon depletion of FfmA as we observed above. The *A. fumigatus AFUA_1g14750* gene is detected by ChIP-seq using either the FLAG-tagged or the wild-type FfmA protein and showed significant induction upon doxycycline-mediated loss of FfmA. This gene encodes the *A. fumigatus* homologue of the *S. cerevisiae* Sfp1 protein that, when overproduced, led to the induction of ribosomal protein gene transcription (Fingerman *et al.* 2003). *AFUA_1g14750* exhibited a log2 of 0.76 and was just below the threshold of 1 we used in the doxycycline-repressed gene expression cutoff. In the case of the *ffmAΔ* strain previously analyzed (Liu *et al.* 2021), this gene did not exhibit significant induction, again illustrating the differences in the expression profile seen when FfmA was acutely depleted versus chronically removed.

The interaction of FfmA with DNA is not likely to be as relatively simple as for AtrR. Our efforts to detect direct DNA-binding of recombinant FfmA centered on use of the promoter regions from the *ffmA* gene itself, *abcG1* and *atrR* (data not shown). While these genes all contain peaks for FfmA binding by ChIP-seq, it is possible that these cannot be recognized by FfmA in vitro. Given that only a fraction of FfmA can interact with AtrR in the cell, binding of this C2H2 protein might require additional interaction partners.

While there are many examples of C2H2 domain-containing proteins functioning as upstream acting sequence specific binding proteins, other functions of this domain are known. A factor from *Drosophila melanogaster* called GAGA factor or GAF has a single C2H2 domain and can bind DNA in vitro but exhibits more complex interactions to determine its DNA binding pattern in vivo including cooperative binding and recruitment of chromatin remodelers (Tang *et al.* 2022). GAF is an example of a pioneer factor that produces changes in nucleosome patterns after binding to its target genes. The control of chromatin structure exerted by GAF after association with DNA is a potential model for the action of FfmA in vivo.

The model that FfmA may act to globally impact gene expression is supported by at least two different observations. First, cells lacking the *atrR* gene have modest phenotypes unless challenged with azole drugs in vitro (Hagiwara *et al.* 2017; Paul *et al.* 2019). This is not the case for *ffmAΔ* null mutants that are severely compromised in growth. Second, doxycycline-mediated depletion of FfmA using the DO-*ffmA* allele caused a dramatic induction of the *has* regulon. Increased expression of these proteins has been shown to cause increased production of the hexadehydroastechrome iron chelator (Yin *et al.* 2013). This chelator is involved in the response to elevated intracellular iron and elicits production of an orange pigment, a phenotype that aided in our linkage between lowered levels of FfmA and *has* regulon expression. Control of the *has* regulon also demonstrated that, while clear AtrR DNA-binding could be seen in the intergenic *hasA/B* promoter, no significant FfmA DNA-binding was observed (Figure 7A). We suggest that loss of FfmA function may indirectly cause stimulation of expression of the has regulon, potentially due to AtrR overexpression as we discovered earlier (Figure 5B). This close relationship between FfmA and AtrR indicates a regulatory connection that was previously unknown. We suggest that FfmA may be a generally-acting chromatin-associated protein, like the Set1 methyltransferase, and the loss of FfmA may alter chromatin structure across the genome. This generalized disturbance could explain the profound growth defect caused by FfmA depletion as well as its impact on expression of AtrR target genes like *abcG1*. An important future goal is to probe the overall chromatin structure of strains with lowered FfmA levels to test this idea.

## Acknowledgements

We thank Drs. Paul Bowyer and Michael Bromley as well as members of the Manchester Fungal Infection Group for many helpful discussions. We thank Dr. Zach Lewis for pointing out the similarity between FfmA and Mot3. This work was supported by NIH AI143198.

## Supplementary Tables

Supplementary table 1. **Significantly differentially regulated genes in dox-off-*ffmA* cells in the absence of doxycycline.** Genes exhibiting at least a log2=1 change and showing P value of <0.05 in DO-*ffmA* cells grown in the absence of doxycycline compared to wild-type are shown.

Supplementary table 2. **Funcat classification of genes showing significant enrichment using FungiFun2 from DO-*ffmA* cells grown without doxycycline**. Enriched genes were only found in the group of log2=-1 genes in DO-*ffmA* cells grown in the absence of doxycycline.

Supplementary table 3. **Significantly differentially regulated genes in dox-off-ffmA cells in the presence of doxycycline.** Genes exhibiting at least a log2=1 change and showing P value of <0.05 in DO-*ffmA* cells grown in the presence of doxycycline compared to wild-type are shown

Supplementary table 4. **Funcat classification of genes showing significant enrichment using FungiFun2 from DO-*ffmA* cells grown with doxycycline**. Enriched genes found with log2=-1 genes in DO-*ffmA* cells grown in the presence of doxycycline.

Supplementary table 5. **Summary of genes that change similarly or differently between *ffmAΔ* or DO-*ffmA*+doxycycline conditions.** Genes that significantly change in data strains lacking ffmA or in the DO-ffmA strain grown in the presence of doxycycline are summarized.

Supplementary table 6. **ChIP-seq peaks common to ChIP reactions performed with either anti-FfmA or anti-FLAG antisera.** ChIP-seq peaks were collected from reactions on wild-type strains using anti-FfmA antibody or on FLAG-FfmA-expressing strain using anti-FLAG antibody. The union of these two data sets is reported.

Supplementary table 7. **Direct target genes enriched via Funcat term analysis using FungiFun2.** Genes detected as common FfmA targets in ChIP-seq data above and also induced in DO-ffmA cells treated with doxycycline are listed.

Supplementary table 8. **Transcriptional proteins enriched via GO term analysis using FungiDB.** Genes detected as common FfmA targets in ChIP-seq data above and also as AtrR targets were analyzed using FungiDB and the transcription factors enriched are listed.

Supplementary table 9. **Asexual sporulation-related proteins enriched via GO term analysis using FungiDB.** Genes detected as common FfmA targets in ChIP-seq data above and also as AtrR targets were analyzed using FungiDB and the genes involved in asexual sporulation are listed.

